# Mechanistic Insights Revealed by YbtPQ in the Occluded State

**DOI:** 10.1101/2021.06.15.448544

**Authors:** Wenxin Hu, Chance Parkinson, Hongjin Zheng

## Abstract

Recently, several ATP-binding cassette (ABC) importers have been found to adopt the typical fold of type IV ABC exporters. Presumably, these importers would function under the transport scheme of “alternating access” like those exporters: cycling through conformations of inward-open, occluded, and outward-open. Understanding how the exporter-like importers move substrates in the opposite direction requires structural studies in all the major conformations. To shed light on that, here we report the structure of yersiniabactin importer YbtPQ from uropathogenic *Escherichia coli* in the occluded conformation trapped by ADP-vanadate (ADP.Vi) at 3.1Å resolution determined by cryo electron microscopy. The structure shows unusual local rearrangements in multiple helices and loops in its transmembrane domains (TMDs). In addition, the dimerization of nucleotide-binding domains (NBDs) promoted by the vanadate trapping is highlighted by the “screwdriver” action happening at one of the two hinge points. These structural observations are rare and thus provide valuable information to understand the structural plasticity of the exporter-like ABC importers.

## Introduction

Although ABC transporters have been extensively studied, new knowledge about these proteins is gained every day. Recently, to everyone’s surprise, three bacterial importers have been reported to adopt the typical fold of type IV ABC exporters ^1^, deviating from all known ABC importers. These importers are yersiniabactin (Ybt) importer YbtPQ from uropathogenic *Escherichia coli* (UPEC) ^2^, as well as IrtAB and Rv1819c from *Mycobacterium tuberculosis* ^3,4^. Such studies have raised immediate interest in understanding how these importers function similarly/differently compared to the known type IV ABC exporters. To do that, structural studies of these exporter-like importers in all their major physiological conformations are necessary. Here, we have continued our study on YbtPQ and expected to shed more light on the topic.

Originally discovered in *Yersinia enterocolitica* ^5^, Ybt is a virulence-associated siderophore molecule synthesized in many pathogens, such as UPEC. UPEC is the main cause of urinal tract infections (UTIs) that affect about 150 million people worldwide each year ^6,7^. During the infections, UPEC secretes multiple siderophores including Ybt. Ybt is able to recognize and chelate various physiologically relevant metal ions, such as Fe^3+^, Cu^2+^, Co^2+^, Ni^2+^, Ga^3+^ and Cr^3+^ ^8^. The metal chelated Ybt is then imported back into the bacteria through outer membrane importer FyuA ^9^ and inner membrane ABC importer YbtPQ ^10^. This process of siderophore mediated metal uptake is one the most prevalent and vital biological processes in microbes ^11,12^. Specifically, in UPEC, deleting genes encoding YbtPQ importer has detrimental effect on the Ybt-mediated metal uptake and later leads to profoundly decreased UTIs in a cystitis mouse model ^13^. Previously, we have determined high-resolution structure of YbtPQ in the inward-open conformation in both the apo and substrate-bound states. The structures have not only revealed its exporter-like fold, but also provided initial information about the substate binding and releasing.

In this study, we successfully trapped YbtPQ in the occluded state using ADP.Vi in the lipid nanodisc environment. We then determined the structure of the occluded YbtPQ to 3.1Å resolution using cryo-electron microscopy single particle reconstruction. In the TMDs, the structure shows a closed periplasmic gate of the translocation pathway similar to that in the inward-open conformation. However, there are considerable conformational changes in at least five transmembrane helices (TMs) and two loops of the TMDs. In the meantime, the cytosolic gate of the pathway is closed by three groups of residues via strong interactions. The NBDs in the occluded state have sharp experimental densities for the two ADP.Vi molecules and are further dimerized compared to those in the inward-open state. Interestingly, this further dimerization is achieved by a ∼180° rotation action of the C-terminal YbtQ helix coupled with lateral movement, acting like a screwdriver. As far as the authors know, such “screwdriver” mode of NBD dimerization has never been reported for any ABC transporters.

## Results and Discussions

### Structure determination of YbtPQ in the occluded conformation

To better understand the dynamics of YbtPQ, it is necessary to reconstruct its entire transport cycle. Previously published YbtPQ structures with (PDB: 6P6J) and without (PDB:6P6I) substrate Ybt-Fe^3+^ are both in the inward-open conformation ^2^. Here, we expected to trap YbtPQ in other conformations using orthovanadate, as it had been demonstrated to be an effective ATP analog to trap various ATPases, such as myosin ^14^, dynein ^15^, as well as multiple ABC transporters ^16–18^. To do that, we used YbtPQ incorporated in the lipid nanodiscs with *E. coli* polar extract lipids (YbtPQ-nanodisc) because this sample was shown to be functional in our previous study ^2^. After vanadate trapping, the ATPase activity of YbtPQ-nanodisc dropped ∼90% (**Figure S1**). Then, we used this sample for single particle cryo-EM analysis and obtained a final reconstruction at 3.1Å resolution (**Figure S2 & 1A, Table S1**). The map has a size of 120Å × 75Å × 60Å, smaller compared to the inward-open YbtPQ structures with a size of 120Å × 90Å × 60Å. The smaller size reflects the strong dimerization between the NBDs, which also makes the cytosolic opening disappear (**Figure 1B**). Analyzed by the web tool MoleOnline ^19^, the translocation pathway in YbtPQ is completely closed, and the central cavity is too small to fit a Ybt-Fe^3+^ substrate. Thus, this map most likely represents the occluded conformation of YbtPQ, a high-energy intermediate state after ATP binding.

**Figure 1.**
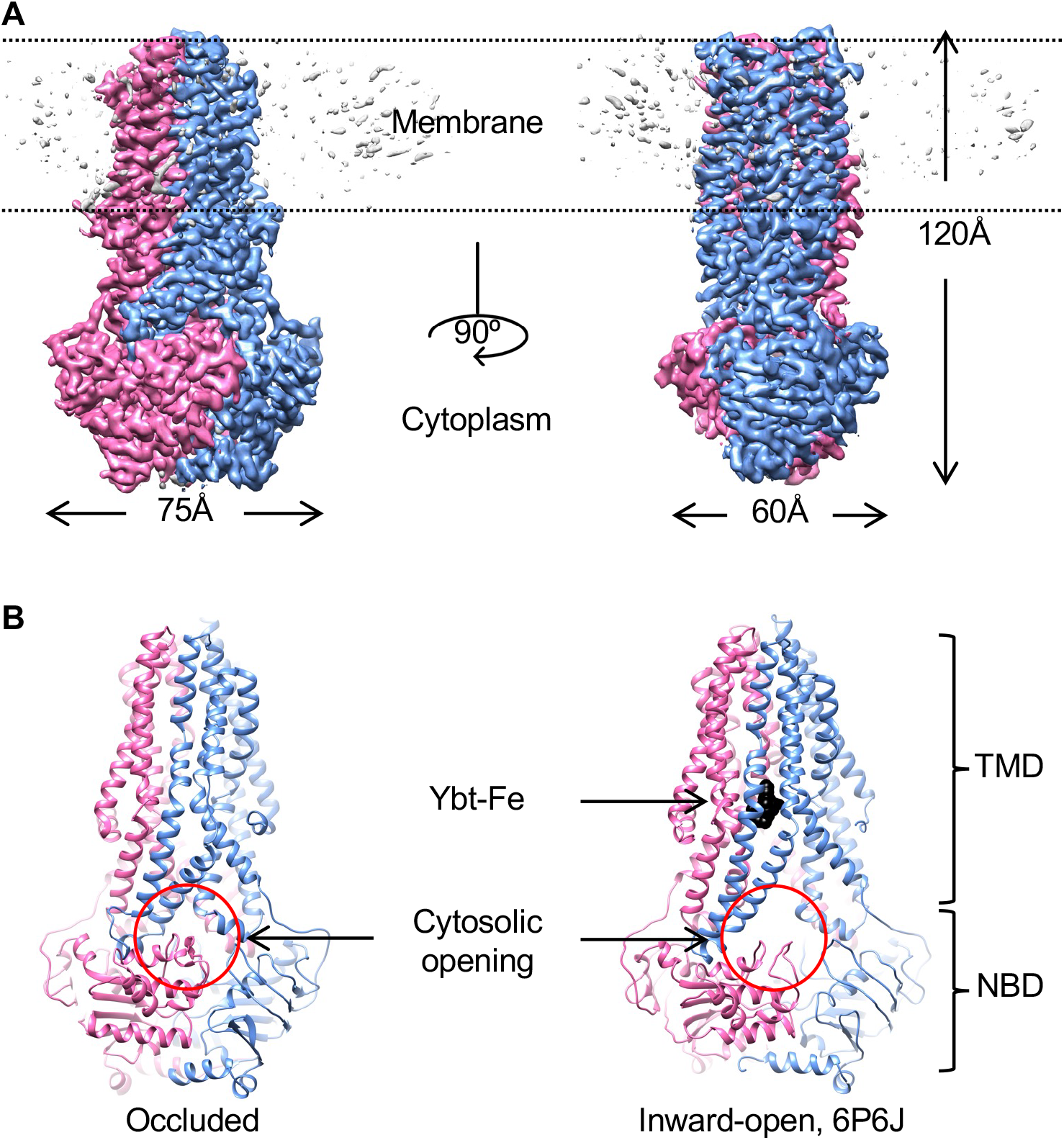
YbtPQ in the occluded state. **A,** cryo-EM map of the ADP-vanadate trapped YbtPQ-nanodisc at 3.1Å resolution. YbtP is in blue, YbtQ is in pink, disordered lipids in the nanodisc is in gray. **B**, The cytosolic opening (red circle) to the substate-binding pocket is closed in the occluded state (left), while open in the inward-open YbtPQ state (right, 6P6J). Substrate Ybt-Fe3+ is shown in black.

### Structural rearrangements of helices and loops in the TMDs

Generally, ABC transporters with the type IV exporter fold function via the alternating access mechanism that the substrate-binding pocket is alternatively exposed in different conformation characterized by the rigid body movement of the TMDs ^20,21^. In YbtPQ, TMD1 is formed by four TMs from YbtP (TM1∼3, TM6) and 2 TMs from YbtQ (TM4, TM5), while TMD2 is formed by four TMs from YbtQ (TM1∼3, TM6) and 2 TMs from YbtP (TM4, TM5), a typical swap of helices in type IV ABC exporters. To reveal the structural differences in the TMDs between inward-open and occluded YbtPQ, we carefully compared those structures. On the periplasmic side, the tip positions of all 12 TMs are the same in both conformations (**Figure 2A**). The periplasmic gate is closed by strong hydrogen bonding among three residues from both TM6: YbtQ-R296, YbtP-E297, and the backbone of YbtP-L289 (**Figure 2A**). Interestingly, the TMDs in YbtPQ do not simply act as two rigid bodies when switching between different conformations, as there are obvious local conformational rearrangements in the TMs and related loops (**Figure 2, S3 & S4**). Specifically, the middle of the following helices: YbtP-TM6, YbtQ-TM6 and YbtQ-TM4-5, are completely twisted with both ends well aligned. The N-terminal part of YbtQ-TM3 connected to the intracellular loop 1 (coupling helix 1 between the TMDs and NBDs, ICL1) moves significantly with part of the helix relaxed. In addition, the loops connecting TM5 and ICL2 (coupling helix 2 between the TMDs and NBDs) in both YbtP and YbtQ are apparently rearranged.

**Figure 2.**
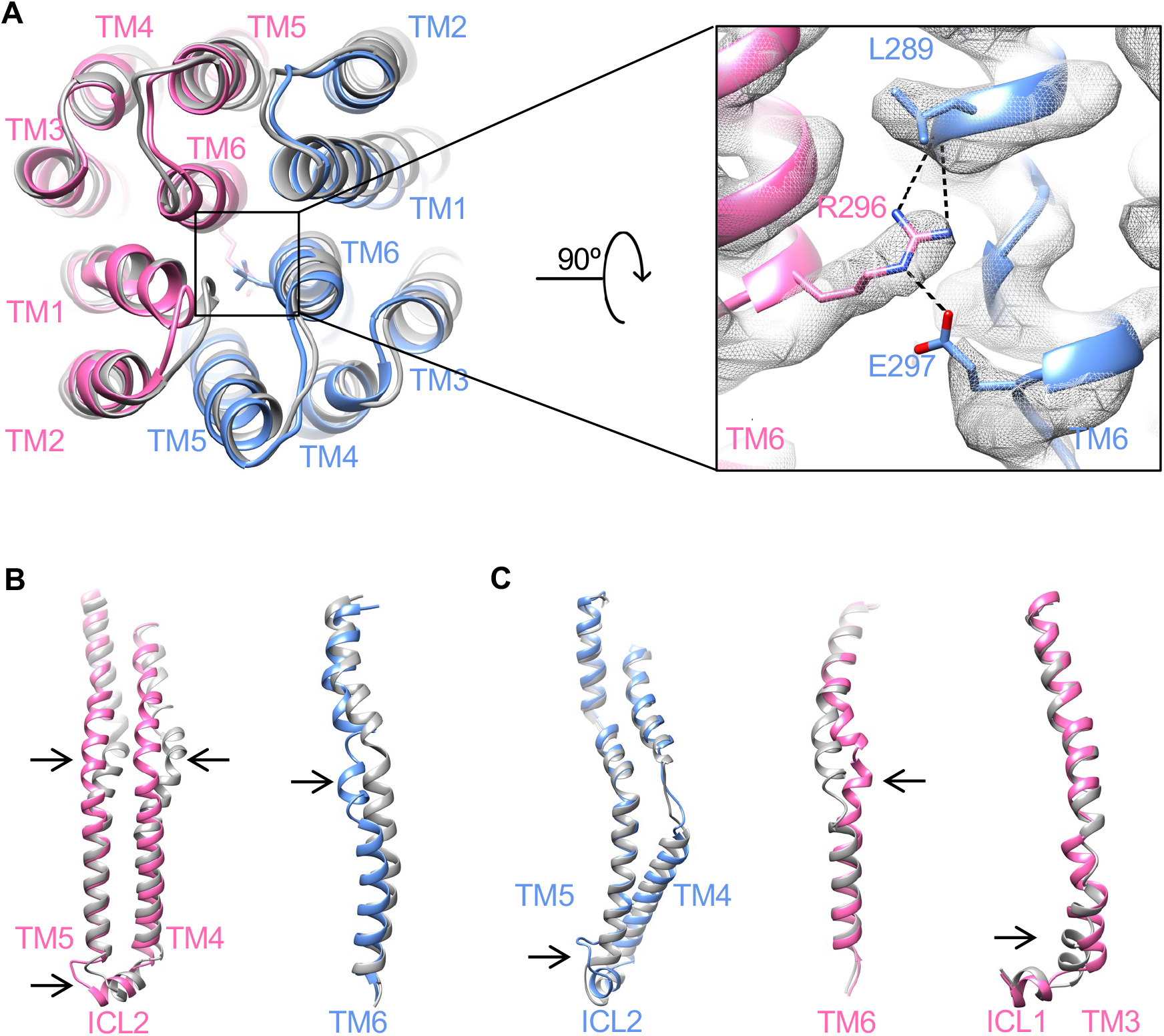
Structural rearrangements in the TMDs during conformational change. **A**. The periplasmic side of individual TMs in the TMDs remains almost identical in both occluded (colored as before) and inward-open (gray, 6P6I) conformations. Residues responsible for the periplasmic gate are highlighted with their experimental densities. **B & C.** TMs and loops from TMD1 (**B**) and TMD2 (**C**) with apparent conformational rearrangements (indicated by the arrows) are shown.

In our opinion, it is reasonable to postulate that the structural rearrangements happening at the end of some helices and loops are necessary to provide flexibility for any conformational changes in a protein. However, it is rather difficult to explain why so many helices in YbtPQ twist only their middle parts. In the literature, one of the closest structural homologs to YbtPQ with different conformations available is the heterodimeric multidrug exporter TmrAB from *Thermus thermophilus* ^18,22^. In TmrAB, during conformational changes between inward-open and occluded states, only one TM, TmrA-TM3, twists its middle part, while other TMs either remain the same or tilt only one end (**Figure S5**). It appears that TMs in YbtPQ are generally much more flexible comparing to those in TmrAB. Thus, in YbtPQ, the flexible TMs could act like springs, making it harder to transmit all the force provided by nucleotide binding/hydrolysis in the NBDs to the periplasmic side of the TMDs. While in TmrAB, the rigid nature of the TMs makes it easier to transmit the force. In consequence, when trapped by ADP.Vi, the conformational switch in YbtPQ is not complete, so it only exists in occluded state. No particles in the outward-open conformation could be found in the 2D classes (**Figure S2**). While in TmrAB, the conformational switch is complete as it exists in both occluded and outward-open states ^18^.

How does YbtPQ close the cytoplasmic side of the translocation pathway in the occluded state? There are three groups of residues responsible for that, all of which are around the circled area (**Figure 1B)**. Specifically, H196 and E200 from YbtP-TM4 interact with K125 from YbtP-TM3 and R318 from YbtP-TM6, respectively (**Figure 3A**). S207 and D209 from YbtQ-TM4 interact with G125 and R123 from YbtQ-TM3, respectively (**Figure 3B**). Q211 from YbtP-TM4 forms strong hydrogen bonding with Q215 from YbtQ-TM4 (**Figure 3C**). Together, these residues help YbtPQ remain in the occluded state.

**Figure 3.**
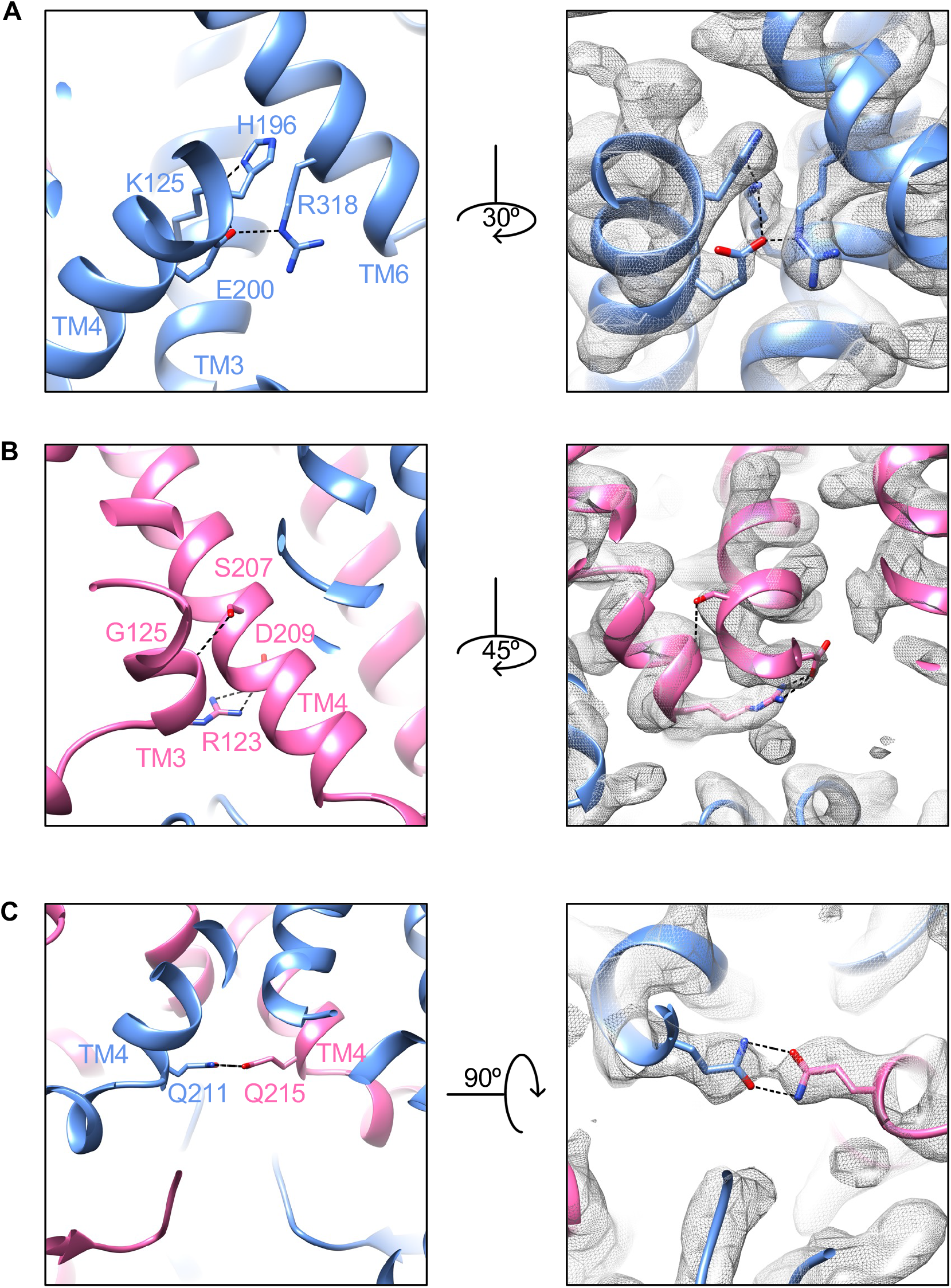
Closure of the cytoplasmic opening mediated by three groups of residues. **A**. Residues at the cytoplasmic side of the substrate-binding pocket from YbtP. **B.** Residues at the back of the opening from YbtQ. **C.** Residues directly bridge TM4s from YbtP and YbtQ.

### Enhanced dimerization of the NBDs by ADP.Vi

How do ADP.Vi molecules interact with the NBDs in YbtPQ? To answer that, we carefully examined the cryo-EM reconstruction and found two clear densities of ADP.Vi in the ATP-binding sites formed in between the two NBDs (**Figure S6 & 4A**). ADP.Vi molecules form canonical interactions with important residues from highly conserved loops and motifs. Specifically, adenines from the two ADP are oriented by the aromatic YbtQ-Y353 and YbtP-Y351 from A-loops via strong π-π interactions. The ribose groups form hydrogen bonds with YbtP-Q485 and YbtQ-E486 from C-loops (ABC signature motifs). The phosphate groups interact with multiple residues from P-loops (Walker A) and C-loops. All these interactions effectively bring the two NBD domains together, so the Q-loops and H-loops are close enough to catalyze the ATP hydrolysis. Here, the orthovanadate molecules, mimicking the hydrolyzed phosphate groups, are specifically stabilized in the binding sites by residues YbtQ-Q426/YbtQ-H538, as well as YbtP-Q425/YbtP-H537, from the Q- and H- loops.

**Figure 4.**
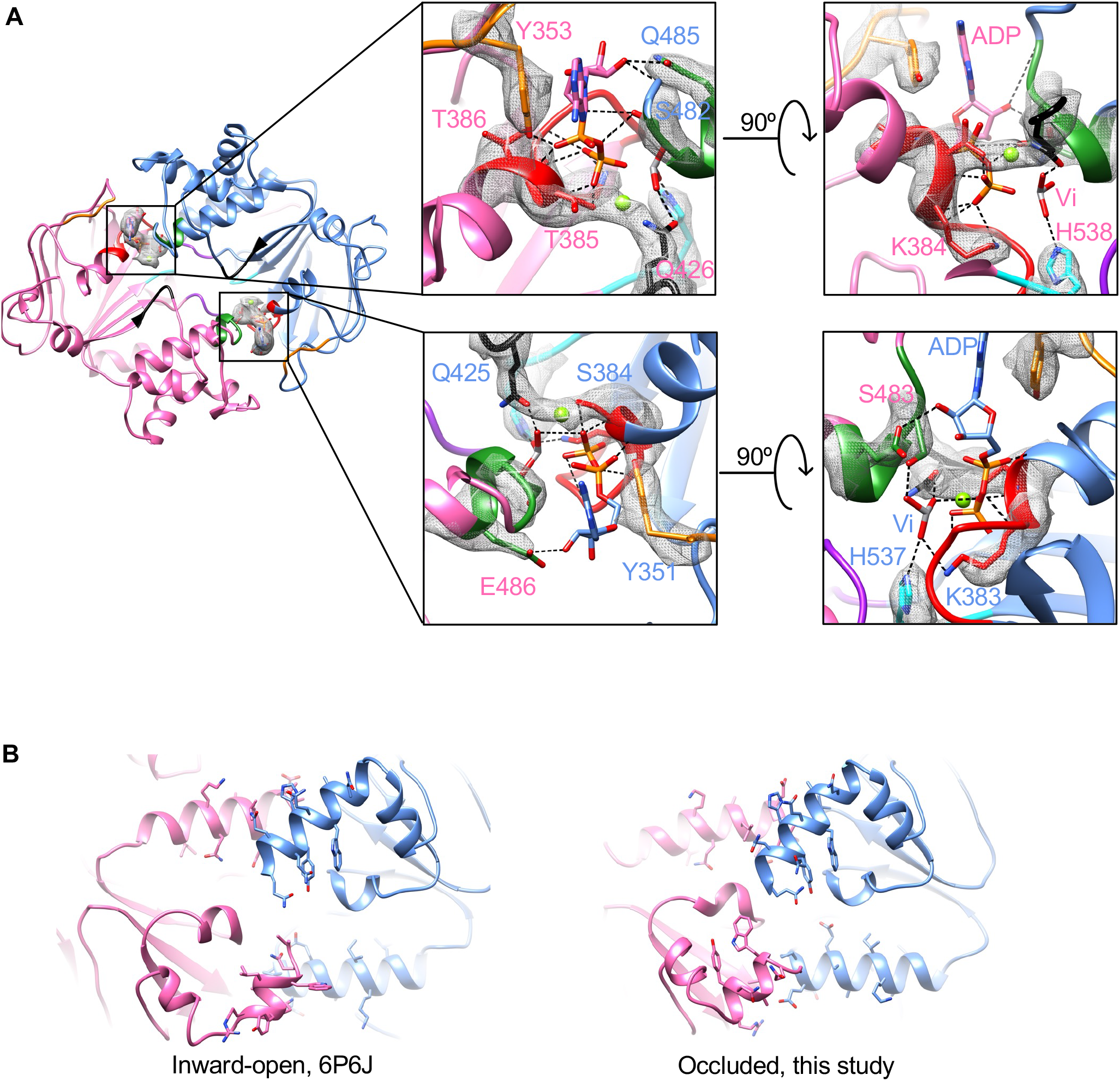
NBD dimerization upon vanadate trapping. **A.** Detailed interactions between ADP.Vi molecules and NBDs. Looking down the translocation pathway from periplasm, the sliced view shows only the NBDs. Conserved P-loop, C-loop, D-loop, A-loop, Q-loop and H-loop are colored as red, green, purple, orange, black and cyan, respectively. Experimental densities of corresponding side chains and ADP.Vi are shown as gray mesh. **B.** Unsymmetrical dimerization mediated by the interactions between C-terminal helix of an NBD and another helix of the opposing NBD. The views shown are the bottom of NBDs in the inward-open and occluded states. Side chains of the residues in those helices are shown in stick mode.

**Figure 5.**
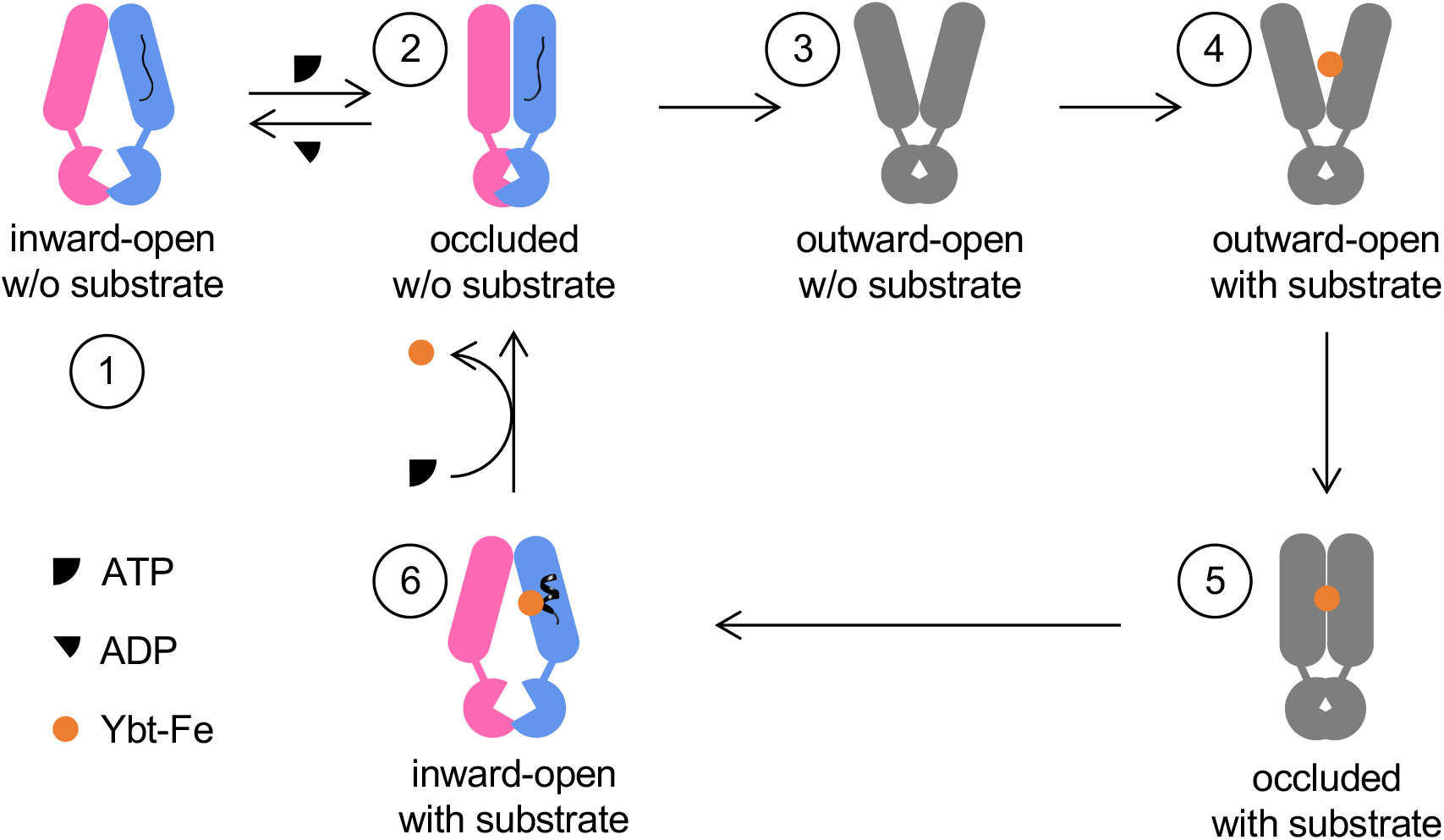
Known YbtPQ structures fit into the proposed import cycle. As discussed in the text, six conformational states of YbtPQ are indicated by the numbers. Colored cartoons represent structures determined in this and previous studies, while gray cartoons represents unknown states.

In the inward-open YbtPQ, we have shown that the NBDs are already dimerized through two sets of helix-helix interactions: C-terminal helix of YbtP interacts with helix I514-M527 in YbtQ, and C-terminal helix of YbtQ interacts with helix E514-L527 in YbtP (**Figure 4B**) ^2^. It is well known that ATP (ADP.Vi in this case) binding promotes further dimerization of the NBDs, as demonstrated in many ABC transporters. This action involves one of the two known types of NBD motion: rocking motion with a pivot at the bottom of the NBDs, and swing-arm motion with a pivot in the TMDs ^23^. To understand the NBD motion in YbtPQ, we morphed YbtPQ between the inward-open and occluded conformations and examined the helix-helix dimerization sites. The result shows that when ADP.Vi binds, the relative positions of C-terminal helix of YbtP and helix I514-M527 in YbtQ do not change too much. The interactions remain largely in between the last 2 visible turns of C-terminal helix of YbtP and the first 2 turn of helix I514-M527 in YbtQ, with slight adjustments of the side chains (**Figure 4B, Movies S1)**. Interestingly, the interactions between C-terminal helix of YbtQ and helix E514-L527 in YbtP are drastically modified. The C-terminal helix rotates ∼180° and moves along the other helix, as if it is a “screwdriver” (**Figure 4B**, **Movies S2**). This type of action, rotating around the helical axis coupled with moving along another helical axis between two interacting helices, has been studied in some cases, such as signal transduction via HAMP domain ^24^ and Rhodopsin ^25^. However, to our knowledge, this kind of unsymmetrical and drastic screwdriver-like action has never been reported in any other ABC transporters.

### Concluding remarks

In this study, we have determined the structure of the exporter-like importer YbtPQ in the occluded state with ADP.Vi bound. Critical residues responsible for the closure of both ends of the substate translocation pathway have been discussed. By comparing the occluded structure to the inward-open structures, we have identified several unique features in YbtPQ. First, TMs in the TMDs are more flexible than other ABC transporters with similar type IV fold, as many of them twist the middle part during the conformational change. Second, NBD dimerization promoted by the vanadate trapping follows a general pivotal rocking motion. However, the two pivot points do not move symmetrically, with one of which goes through the screwdriver-like action.

How does the new structure fit into the whole import cycle? Let’s discuss the cycle in the following six steps. 1, we can start from the inward-open conformation without substrate/ATP bound, whose structure has been determined previously (PDB: 6P6I). 2, considering the high ATP concentration in a living cell, it’s reasonable to suggest that ATP will bind to YbtPQ and try to start the import event. Our new structure is likely a mimic of this state. It seems that ATP binding could only change the protein from inward-open to occluded, but not all the way to outward-open. The reason is probably because TMs in YbtPQ are not rigid enough to transmit all the mechanical force from the NBDs all the way up to its periplasmic gate. It is also possible that the lipids in nanodiscs are restrained by the scaffold protein, or the types of lipid molecules in nanodiscs are not ideal, so the periplasmic side refuses to open. Last but not least is that there may be a periplasmic soluble protein to help YbtPQ switching to outward-open, though previous efforts of looking for such a partner have not been successful ^8,10,26^. Nevertheless, this incomplete conformational switch is beneficial to YbtPQ’s import function, as it is less likely to accidentally export the substrate. At this step, ATP hydrolysis will bring YbtPQ back into the inward-open state, which is the futile ATPase activity observed for almost all ABC transporters. 3, for the reasons not fully understood yet, YbtPQ shall be able to reach the outward-open conformation. 4, substrates will then be recognized and bound to the substrate-binding pocket exposed to the periplasm. 5, YbtPQ changes back to the occluded state with substrate bound. 6, After ATP hydrolysis and ADP released from the NBDs, YbtPQ goes back to the inward-open conformation with substate still bound (PDB: 6P6J). Previously, we proposed that the substrate release should be facilitated by the unwinding of YbtP-TM4, going directly from step 6 to step 1 ^2^. Here, we would like to update this model by adding step 2 into the substrate releasing process. After import (step 6), YbtPQ will bind ATP again to promote conformational change to the occluded state (step 2). This conformational change closes the cytosolic gate, crashes the substrate-binding pocket, and likely to provide energy to unwind the helix YbtP-TM4 (also observed in the occluded structure in **Figure 2B**), so to release the substrate. Then, YbtPQ can release ADP and switch back to step 1.

### Materials and Methods

### Protein expression and Purification

Wild-type full-length YbtPQ importer from uropathogenic *E. coli* UTI89 were over-expressed and purified as previously reported ^2^. Briefly, the heterodimeric YbtPQ with a N-terminal His-tag on YbtP was expressed in *E. coli* strain of BL21(DE3) C43 (Sigma-Aldrich) at 18°C overnight with 0.2mM Isopropyl β-D-1-thiogalactopyranoside (IPTG, UBPBio). Membrane fractions from 3L of harvested cells were prepared in 20mM Tris pH 7.5, 150mM NaCl (Buffer A), 1mM phenylmethylsulfonyl fluoride (PMSF). 1% lauryl maltose neopentyl Glycol (LMNG) was used to solubilize YbtPQ from the membrane. YbtPQ was purified by Ni-NTA affinity column followed by a gel filtration on a Superose 6 column (GE Healthcare Life Sciences) in Buffer A with 0.002% LMNG, and then concentrated to ∼6mg/ml.

### Vanadate trapping of YbtPQ in lipid nanodiscs

Purified YbtPQ was reconstituted into the *E. coli* polar extract lipids (Avanti Polar Lipids) with the membrane scaffold protein MSP1D1 (Addgene) as previous reported ^2^. Briefly, MSP1D1 was expressed at 37°C in *E. coli* BL21(DE3) strain, purified by one-step affinity chromatography using Ni-NTA resin, and concentrated to ∼5mg/ml. The stock of *E. coli* polar extract lipids at 50mM was made in Buffer A with 150mM LMNG. To reconstitute the nanodiscs, purified YbtPQ, MSP1D1 and *E. coli* polar extract lipids were mixed at a molar ratio of 1:8:400 on ice for 30 min before adding 0.2mg Bio-Beads for an overnight incubation at 4°C. The assembled YbtPQ-nanodiscs were further purified by gel filtration on a Superose 6 column in Buffer A. The corresponding peak was pooled together and concentrated to ∼6mg/ml. The stock of 100mM orthovanadate solution was prepared as previously reported ^14^. Purified YbtPQ-nanodiscs were incubated with 20mM vanadate, 18mM ATP and 54mM MgCl2 for 15min at 37°C. 36mM fresh ATP was added to the solution and sit at room temperature for 5 min. The sample was finally purified by gel filtration on a Superose 6 column in Buffer A.

### ATPase activity assay

The ATPase activity was determined as previously reported ^2^, using an ATPase/GTPase Activity Assay Kit (Sigma-Aldrich) featuring a detectable fluorescent product from malachite green reacting with released phosphate group. The bar graph of the ATPase activities of YbtPQ-nanodisc with and without vanadate trapping were prepared in Excel. The experiments were repeated three times independently.

### Cryo-EM sample preparation and data acquisition

3ul of vanadate-trapped YbtPQ-nanodiscs at ∼2mg/ml were applied to plasma-cleaned C-flat holy carbon grids (1.2/1.3, 400 mesh, Electron Microscopy Sciences) and prepared using a Vitrobot Mark IV (Thermo Fisher Scientific) with the environmental chamber set at 100% humidity and 4°C. The grids were blotted for 3.5∼4.5s and then flash frozen in liquid ethane cooled by liquid nitrogen. Cryo-EM data was collected on a Titan Krios (Thermo Fisher Scientific) operated at 300keV and equipped with a K3 direct detector (Gatan). 5,152 movies were recorded with a calibrated pixel size of 0.83Å, defocus range of −1 ∼ −2.5μm, 50 frames with a total dose of ∼60 electrons/Å^2^.

### Cryo-EM data processing, model building, refinement, and validation

The data was processed using the software suit cryoSPARC v2.15.0 ^27^. Briefly, the movies were motion corrected using Patch motion correction, and their Contrast transfer function (CTF) parameters were estimated using Patch CTF. Micrographs with a calculated defocus range of −0.8μm ∼ 2.8μm, fitted resolution of better than 8Å, as well as global motion of less than 35 pixels were selected for further processing. A total of ∼2 million particles were picked using Blob picker with the minimum diameter set to 60Å and the maximum particle diameter set to 150Å. These particles were extracted with a box size of 320 × 320 pixels and subjected to two-rounds of 2D classification. ∼900,000 particles were selected for the ab-initio reconstruction of 4∼6 initial models. These initial models were heterogeneously refined, and the best group with 340,080 particles was selected for further homogeneous refinement. The final reconstruction was estimated to be at 3.07Å resolution based on the gold-standard ^28^ Fourier shell correlation (FSC) with a cut-off of 0.143. Local resolution estimation was done in cryoSPARC. The model building was carried out in Coot program ^29^, guided by the previously published structure of YbtPQ in the inward-open conformation (PDB: 6P6J). The final rounds of model refinement were carried out by real space refinement in PHENIX ^30^ with secondary structure restraints imposed. The quality of the model was assessed by MolProbity ^30^. To validate the refinement, the model was refined against half-maps, and FSC curves were calculated. The final statistics was shown in the Extended Data Table 1. All figures were prepared using the programs UCSF Chimera ^31^.

## Acknowledgements

A portion of this research was supported by NIH grant U24GM129547 and performed at the Pacific Northwest Center for Cryo-EM (PNCC) at Oregon Health & Sciences University (OHSU) and accessed through EMSL (grid.436923.9), a DOE Office of Science User Facility sponsored by the Office of Biological and Environmental Research. Some of the work was also performed at the Stanford-SLAC Cryo-EM Center (S2C2) (supported by the NIH U24 GM129541). Specifically, we thank Mr. Theo Humphreys from PNCC for the final Cryo-EM data collection.

## Author contributions

Hu, W., Parkinson, C. and Zheng, H. designed the study, performed the experiments, analyzed the data, and wrote the manuscript.

## Data and material availability

Cryo-EM map of YbtPQ was deposited in the Electron Microscopy Data Bank under the accession code EMD-24234. Coordinates of the atomic model of YbtPQ was deposited in the Protein Data Bank under the accession code 7N8A. All other data are available from the corresponding author upon reasonable request.

---

**Figure S1.**
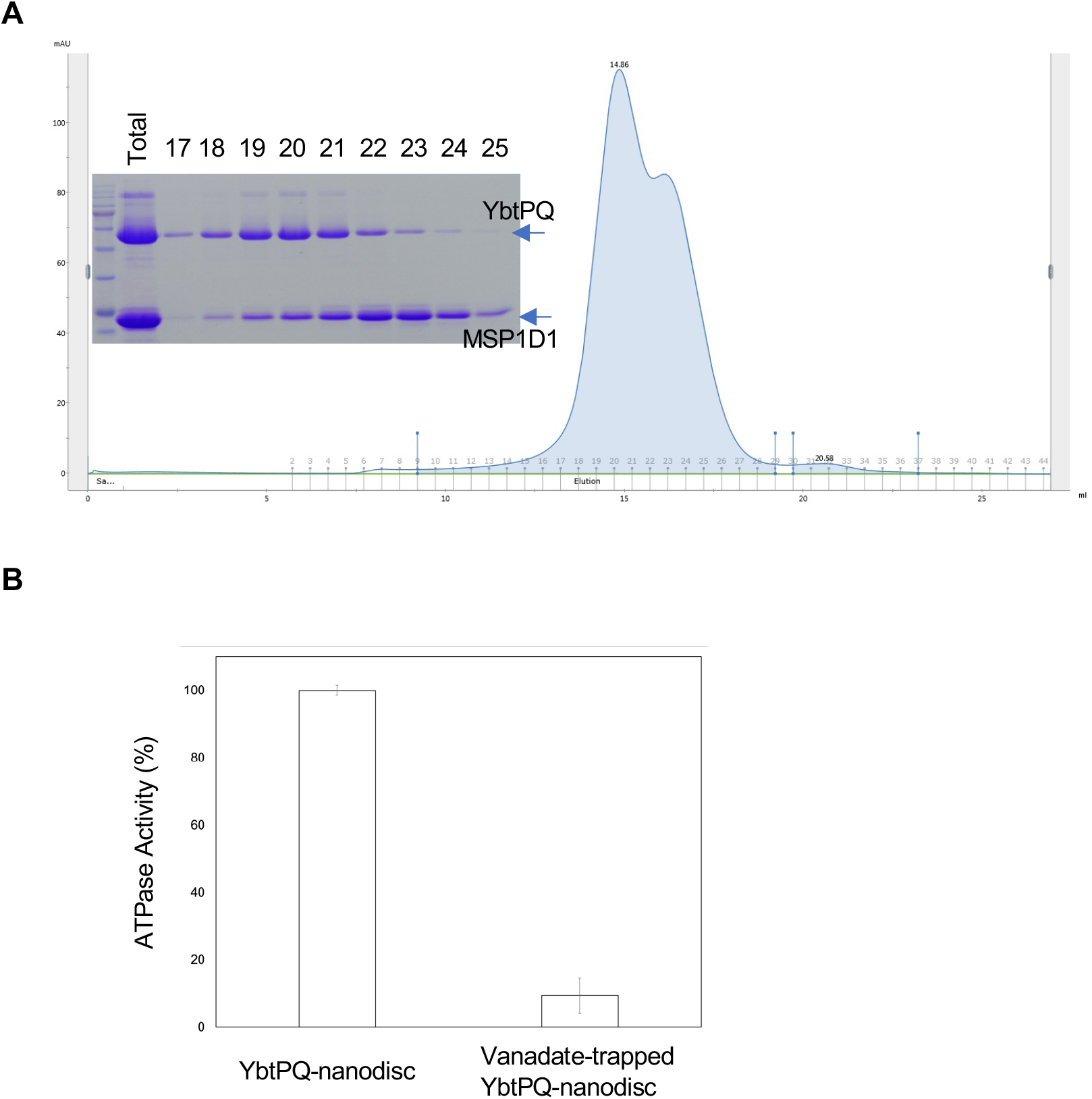
YbtPQ-nanodisc used for cryo-EM analysis. **A,** Gel filtration profile of the YbtPQ-nanodisc assembly and SDS-PAGE analysis of corresponding elution fractions. The first peak (fractions 18∼21) was collected and subjected to vanadate trapping. **B**, ATPase activity of vanadate-trapped YbtPQ-nanodisc dropped ∼90% comparing to non-trapped YbtPQ-nanodisc.

**Figure S2.**
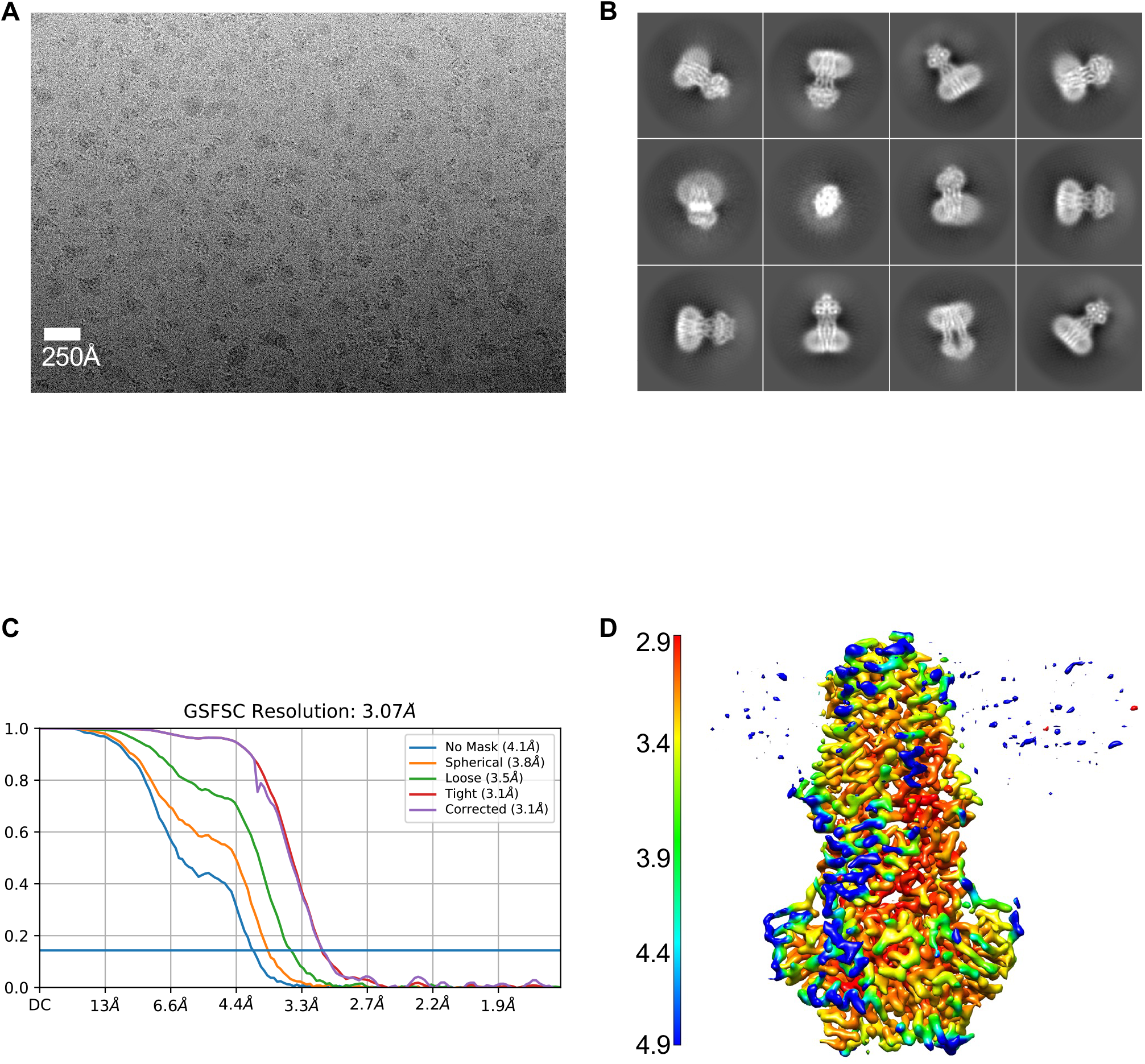
Results of cryo-EM analysis of vanadate-trapped YbtPQ-nanodiscs. **A**, A representative cryo image. **B**, Selected 2D averages. **C**, Gold-standard FSC curve. **D**, Final YbtPQ map colored by local resolution estimation.

**Figure S3.**
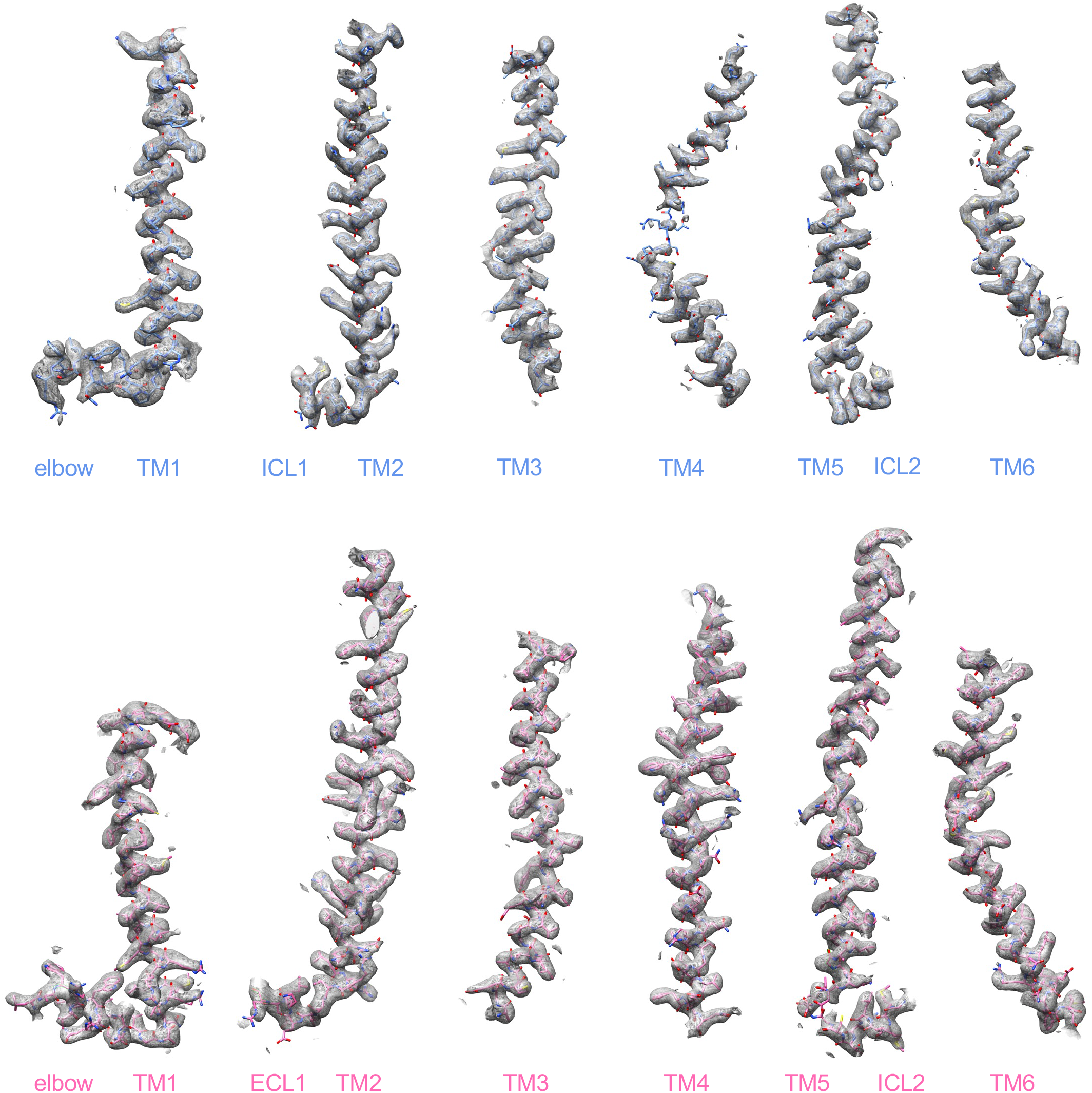
Quality of the experimental densities in the TMDs. All 12 TMs, elbow helices and ICLs are shown. YbtPQ is colored as before.

**Figure S4.**
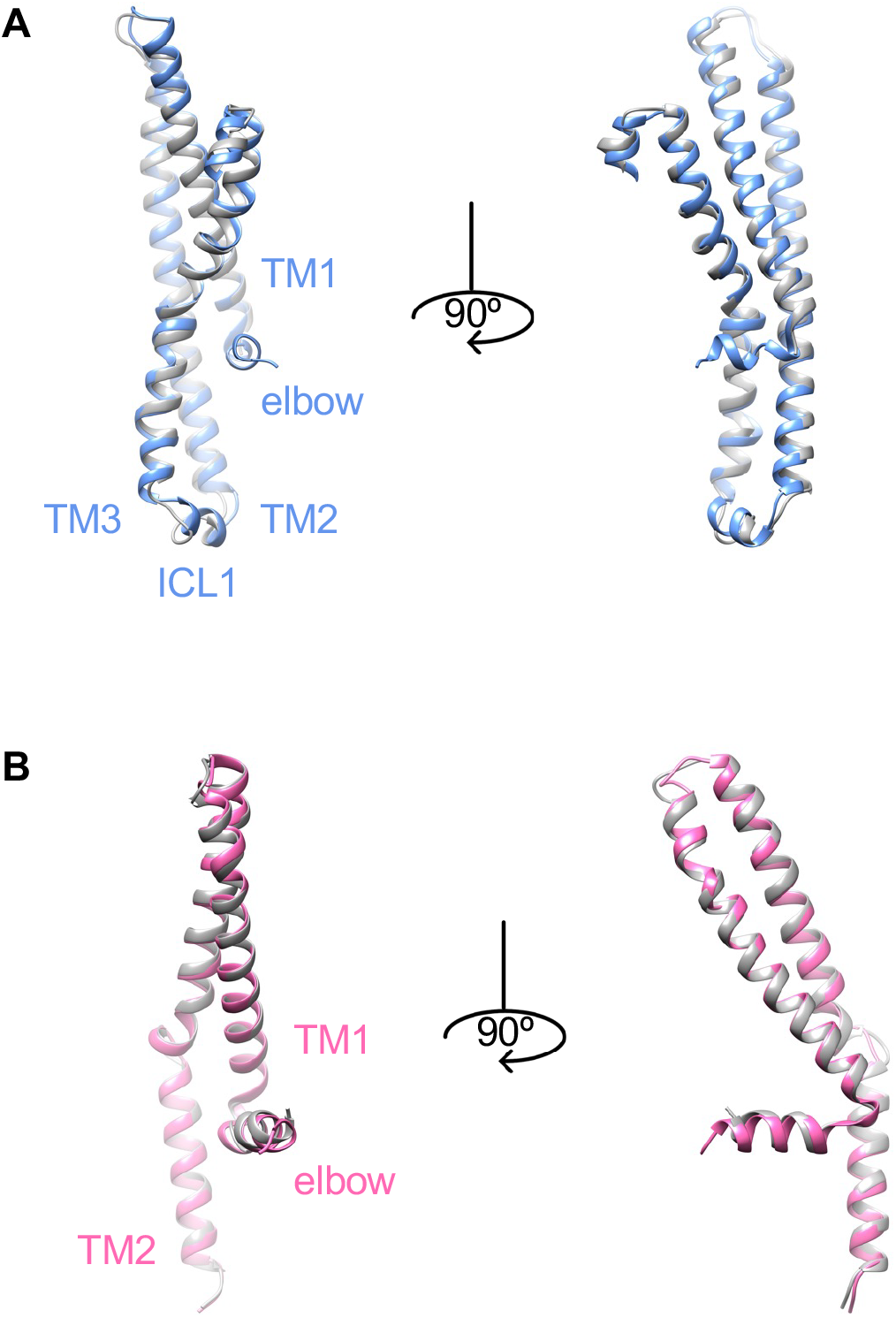
TMs in YbtPQ without apparent structural rearrangement. **A**. TM1-3 in YbtP. **B**. TM1-2 in YbtQ. Occluded YbtPQ is colored as before, while inward-open YbtPQ is in gray.

**Figure S5.**
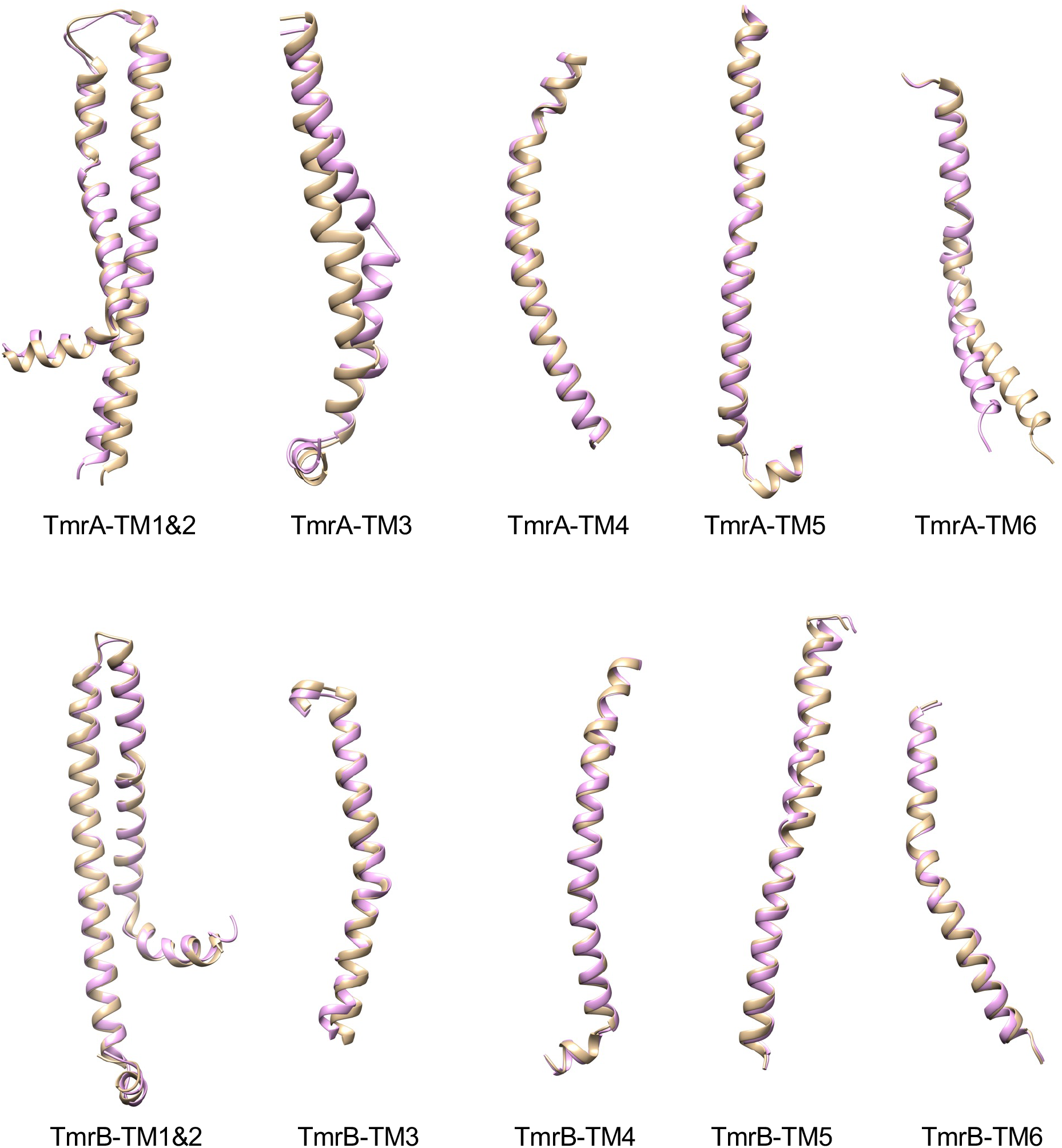
Helical rearrangement in TmrAB. All TMs from Inward-open (6RAF, brown) and occluded (6RAI, pink) states are aligned individually. Only TmrA-TM3 shows similar twist in the middle as in YbtPQ.

**Figure S6.**
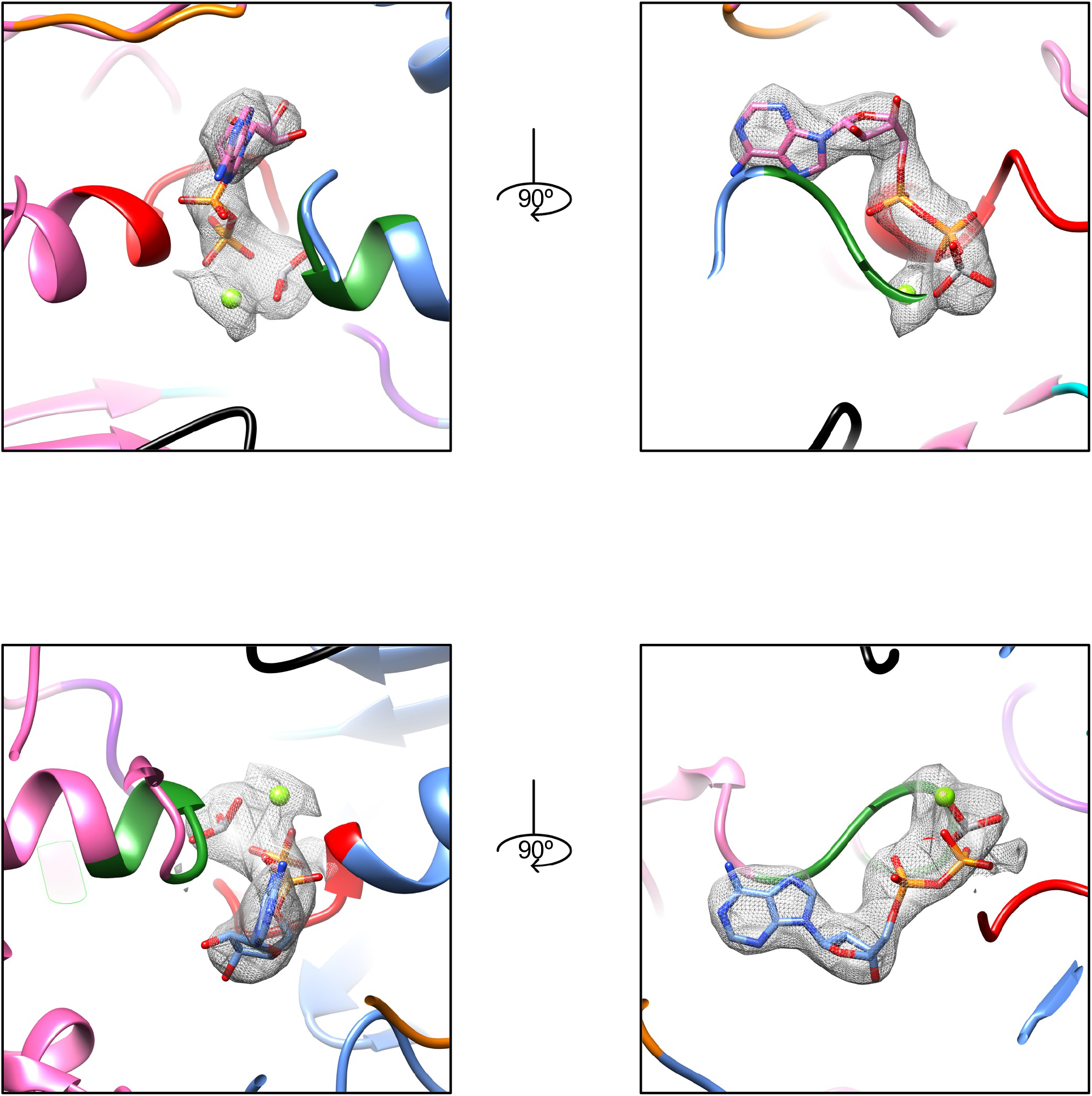
Two experimental densities of ADP-vanadate with the models fitting inside. Density in the top panels is the top-left one in Figure 4A, and the density in the bottom panels is the bottom-right one in Figure 4A.

**Table S1:** Cryo-EM data collection, refinement and validation statistics.

|  |  |
| --- | --- |
|  | YbtPQ |
| **Data Collection and Processing** |  |
| Microscope | Titan Krios |
| Voltage (kV) | 300 |
| Magnification (nominal) | 92,000 |
| Electron Dose (e<sup>-</sup>/Å<sup>2</sup>) | 60 |
| Camera | Gatan K3 |
| Defocus range (um) | -1 ~ -2.5 |
| Pixel size (Å) | 0.83 |
| Movies collected | 5152 |
| Symmetry imposed | C1 |
| Final particle images (no.) | 340,080 |
| Map resolution (Å) | 3.1 |
| FSC cutoff | 0.143 |
| Map resolution range | 2.9 ~ 4.9 |
| Sharpening B-factor (Å<sup>2</sup>) | -105.5 |
| Software used to process data | cryoSPARC |
| **Refinement statistics** |  |
| Initial model used | 6P6I |
| Model resolution (FSC = 0.5) | 2.9 |
| Correlation Coefficient (Mask) | 0.83 |
| Number of protein atoms (non-H) | 8970 |
| Residues | 1150 |
| R.m.s. deviations |  |
| Bonds (Å) | 0.007 |
| Bond angles (°) | 0.781 |
| **Validation** |  |
| MolProbity score | 2.31 |
| Clash score | 33.89 |
| Poor rotamers (%) | 0.00 |
| **Ramachandran plot** |  |
| Favored (%) | 95.77 |
| Allowed (%) | 4.23 |
| Disallowed (%) | 0 |
| EMDB access code | EMD-24234 |
| PDB access code | 7N8A |

